# A recurrence based approach for validating structural variation using long-read sequencing technology

**DOI:** 10.1101/105817

**Authors:** Xuefang Zhao, Alexandra M. Weber, Ryan E. Mills

## Abstract

Although there are numerous algorithms that have been developed to identify structural variation (SVs) in genomic sequences, there is a dearth of approaches that can be used to evaluate their results. The emergence of new sequencing technologies that generate longer sequence reads can, in theory, provide direct evidence for all types of SVs regardless of the length of region through which it spans. However, current efforts to use these data in this manner require the use of large computational resources to assemble these sequences as well as manual inspection of each region. Here, we present VaPoR, a highly efficient algorithm that autonomously validates large SV sets using long read sequencing data. We assess of the performance of VaPoR on both simulated and real SVs and report a high-fidelity rate for various features including overall accuracy, sensitivity of breakpoint precision, and predicted genotype.

## INTRODUCTION

Structural variants (SVs) are one of the major forms of genetic variation in humans and have been revealed to play important roles in various diseases including cancers and neurological disorders (1,2). Various approaches have been developed and applied to paired-end sequencing to detect SVs in whole genomes (3–5), however individual algorithms often exhibit complementary strengths that sometimes lead to disagreements as to the underlying variant. The emergence of long read sequencing technology, eg. Single Molecule Real-Time (SMRT) sequencing from Pacific Biosciences (PacBio), can provide direct evidence for the presence of an SV. Current strategies make use of de novo assembly to create large contigs that can be cross-referenced with a putative SV using manual inspection of the subsequent recurrence (dot) plot (6). These types of dot plots have been used for decades to examine the specific features of sequence alignments (7), however they require manual curation and, coupled with the computational costs of sequence assembly, are time-consuming and inefficient at scale for the high throughput validation of large sets of SVs.

Here, we present a high-speed long read based assessment tool, VaPoR, that scores each SV prediction by autonomously analyzing the recurrence of windows within a local read against the reference genome in both their original and rearranged format per the prediction. A positive score of each read on the altered reference, normalized against the score of the read on the original reference, supports the predicted structure. A baseline model is constructed as well by interrogating the reference sequence against itself at the query location. We show that our approach can quickly and accurately distinguish true from false positive predictions of both simple and complex SVs as well as their underlying genotypes and is also able to assess the breakpoint accuracy of individual algorithms.

## MATERIAL AND METHODS

### VaPoR Workflow

VaPoR takes in aligned sequence reads in BAM format and predicted SVs (>50bp) in various formats including VCF and BED. Evaluation of an SV is performed by comparing long reads that go through the event against reference sequences in two formats: (a) the original human reference to which the sample is aligned and (b) a modified reference sequence altered to match the predicted structural rearrangement. A recurrence matrix is then derived by sliding a fixed-size window with 1bp step through each read to mark positions where the read sequence and reference are identical. The matching patterns are then assessed as to the validity of the SV as described below and a validation score is reported. Given the large variance of SVs lengths, each SV is stratified into one of two groups: smaller SVs that can be completely encompassed within multiple (>10 by default) long sequences and larger events that are rarely covered by individual long reads, with different statistical model applied. The VaPoR workflow is briefly summarized in Figure 1.

**Figure 1.**
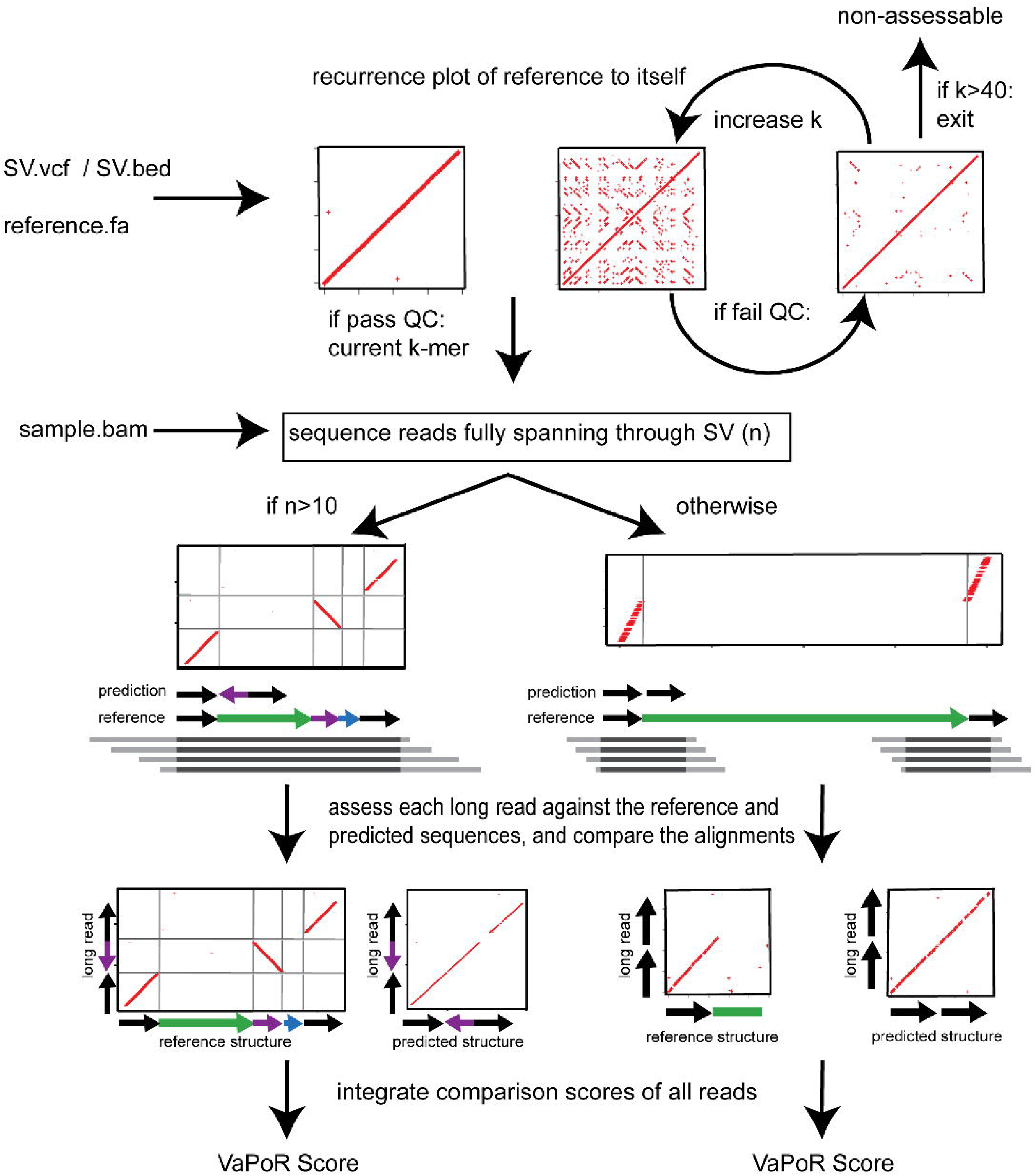
Flowchart describing the VaPoR algorithm. As input, the algorithm requires a set of structural variants in either VCF or BED format, a series of long reads and/or sequence contigs in BAM format, and the corresponding reference sequence. VaPoR then interrogates each variant individually at its corresponding reference location, assesses the quality of the region and assigns a score.

#### Small Variants Assessment

For an SV *k* in sample *s* that is covered by *n* reads, the recurrence matrix between each read and the reference sequences in original (*R_o_*) and altered (*R_a_*) format is calculated. The vertical distance between each record (*X_i,k,s,Rx_*, *Y_i,k,s,Rx_*) in matrix *x* and the diagonal *X_i,k,s,Rx_*, (*X_i,k,s,Rx_*) line is calculated as *d_i,k,s,Rx_* = abs(*X_i,k,s,Rx_* - *Y_i,k,s,Rx_*, and the average distance of all records would be exported as the score of each matrix:

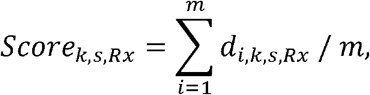

where *m* is the total number of records in the matrix. Sequences that share higher identity with the read shall have a lower *Score_k,s,Rx_,* such that the score of each read is normalized as:

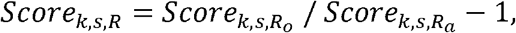

where a positive *Score_k,s,R_* represents the superiority of the predicted structure versus the original and vise versa for negative *Scorer_k,s,R_*, with one exceptional case where there exists a duplicated structure in the predicted SV such that the predicted structure would show higher *Score_k,s,R_* due to the multialignment of duplicated segments. To correct for duplications, VaPoR adopts the directed distance *d_i,k,s,Rx_* = *X_i,k,s,Rx_*, - *Y_i,k,s,Rx_* instead such that the distance contributed by centrosymmetric duplicated segments would offset each other.

#### Large Variants Assessment

For larger SVs where there are few, if any, long reads that can transverse the predicted SV, VaPoR assesses the quality of each predicted junction instead using:

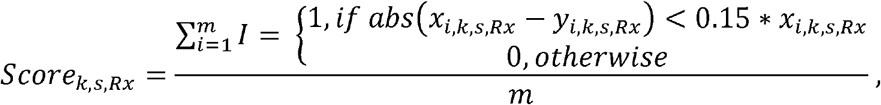

where a larger *Score_k,s,Rx_* represents higher similarity between the read and the reference sequence. The normalized scores of each read is then defined as:

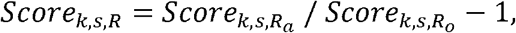

#### VaPoR Score Calculation

With a score assigned to each read spanning through the predicted structural variants, the VaPoR score (*Score_k,s_*) is summarized as:

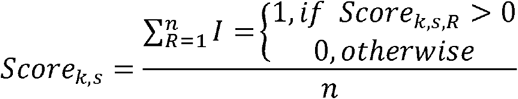

to represent the proportion of long reads supporting predicted structure.

The highest supportive score (max(*Score_k,s,R_*)) is also reported as a reference for users to meet the specific requirement of their study design, for which we recommend 0.1 as the cutoff.

#### Flexible window size

By default, VaPoR uses a window size of 10bp and requires an exact match between sequences, though these can be changed to user-defined parameters. However, many regions of the genome contain repetitive sequences resulting in an abundance of spurious matches in the recurrence matrix, thus introducing bias to the assessment. To address this, VaPoR adopts a quality control step by iteratively assessing the reference sequence against itself and tabulating the proportion of matches along the diagonal. The window size initially starts at 10bp and iteratively increases by 10bp until either (a) the proportion of matches on the diagonal exceeds 40% and the current window size is kept or (b) the window size exceeds 40bp whereby the event will be labeled as ‘non-assessable’ and excluded from the evaluation.

## VaPoR Accuracy Assessment

### Simulated Data

Non-overlapping simple deletions, inversions, insertions and duplications as well as complex structural variants as previously categorized (3) were independently incorporated into GRCh38 in both heterozygous and homozygous states, excluding regions of known difficult regions of the genome as described from the ENCODE project (8). Detailed descriptions of each simulated SV types simulated are summarized in Supplementary Tables 1–3. We applied PBSIM (9) to simulate the modified reference sequences to different read depth ranging from 2X to 70X with parameters difference-ratio 5:75:20. length-mean 12000, accuracy-mean 0.85 and model_qc model_qc_clr. Simulated data can be obtained from https://umich.box.com/v/vapor.

**Table 1.**
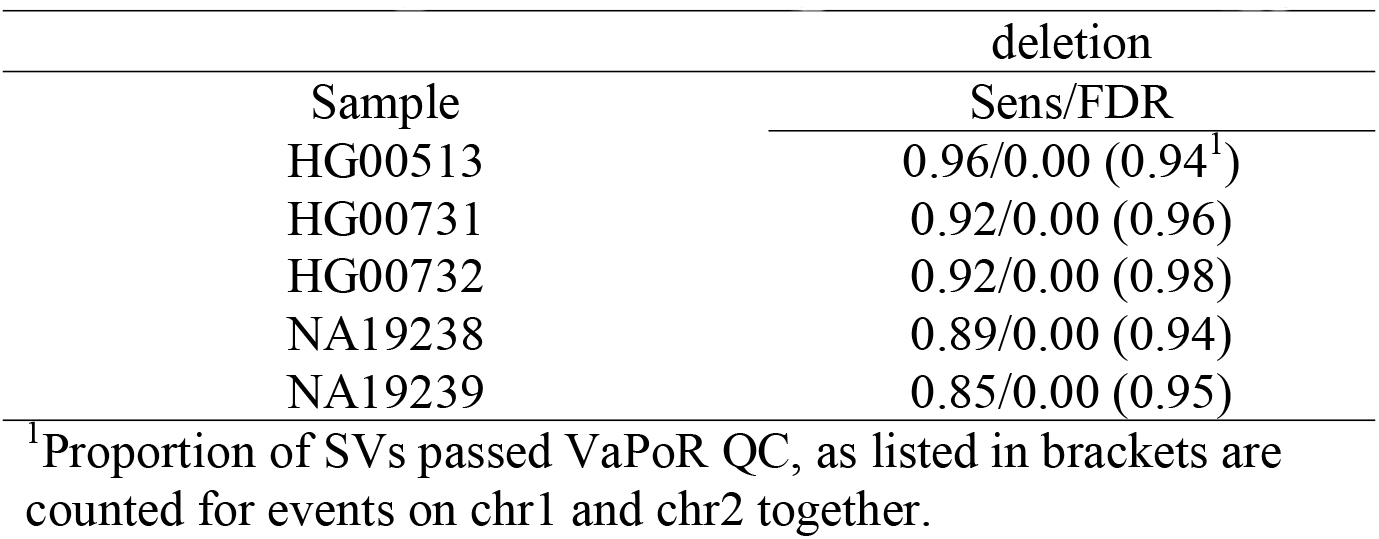
Sensitivity and false discovery rate of different SV types

| Sample | Sens/FDR |
| --- | --- |
| HG00513 | 0.96/0.00 (0.94<sup>1</sup>) |
| HG00731 | 0.92/0.00 (0.96) |
| HG00732 | 0.92/0.00 (0.98) |
| NA19238 | 0.89/0.00 (0.94) |
| NA19239 | 0.85/0.00 (0.95) |

<sup>1</sup>Proportion of SVs passed VaPoR QC, as listed in brackets are counted for events on chr1 and chr2 together.

### Real Data

We also applied VaPoR to a set of diverse samples (HG00513 from CHS, HG00731 and HG00732 from PUR, NA19238 and NA19239 from YRI) that were initially sequenced by the 1000 Genomes Project (1KGP) and for which a high quality set of SVs were reported in the final phase of the project (10). These samples were recently re-sequenced using PacBio and therefore provides a platform for assessing VaPoR on known data. The 1000 Genomes Project Phase 3 data were obtained from ftp://ftp-trace.ncbi.nih.gov/1000genomes/ftp/phase3/integrated_sv_map/ and lifted over to GRCh38. PacBio sequence data were obtained from http://ftp.1000genomes.ebi.ac.uk/vol1/ftp/datacollections/hgsvsvdiscovery/.

## RESULTS

We assessed the performance of VaPoR on both simulated sequences and real genomes from the 1000 Genomes Project to assess the following characteristics: sensitivity and false discovery rate on validating structural variants in simple and complex structures; sensitivity of VaPoR on validating different levels of predicted breakpoint efficacy; stratification of VaPoR scores by genotype; and time and computational cost of VaPoR.

### VaPoR on Simulated Data

We applied VaPoR to simulated simple deletions, inversions, insertions and duplications as well as complex structural variants and first assessed the proportion of SVs that VaPoR is capable of interrogating (i.e. passed VaPoR QC). We found that VaPoR can successfully evaluate >80% of insertions, >85% deletion-duplications and >90% SVs in all other categories when the read depth is 10X or higher. We then assessed the sensitivity and false discovery rate (FRR) at different VaPoR score cutoffs and found that when considering different types of SVs across various read depths of, most of the SV types can achieve a sensitivity >90% with false discovery rate <10% at a VaPoR score cutoff of 0.15 (Supplemental Figures 1–2). We further observed that there were no significant changes of sensitivity or false discovery rate once the read depth was at or above 20X and is consistent across different SV types (Figure 2, Supplemental Tables 1–3).

**Figure 2.**
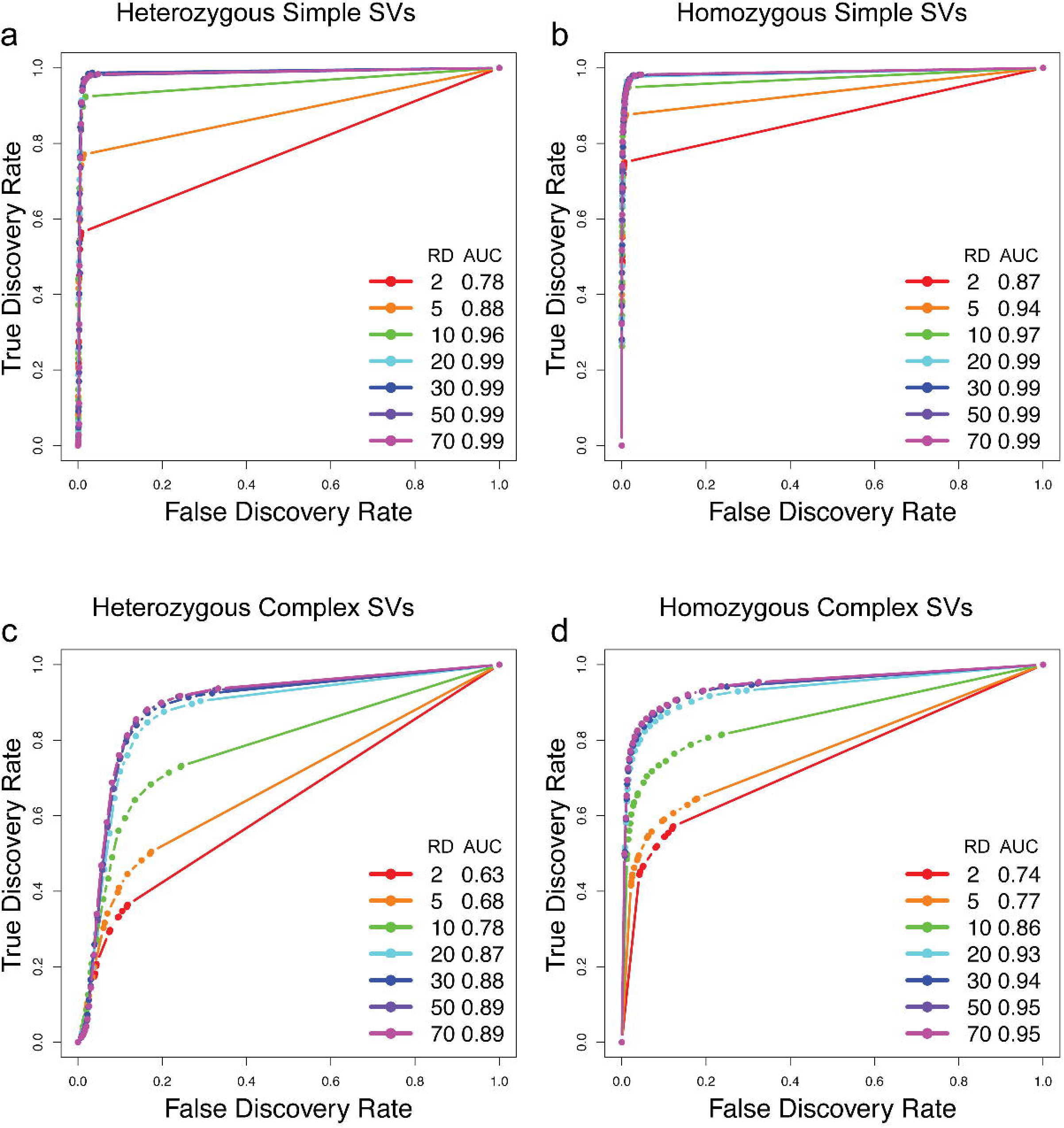
Accuracy of VaPoR on simulated heterozygous and homozygous SVs at varying degrees of sequence coverage and VaPoR score cut-offs. Receiver operator curves (ROC) are shown for simple deletions, duplications and inversions (a,b) as well as complex rearrangements including inverted duplications and deletion-inversion rearrangements (c,d).

### VaPoR on 1000 Genomes Project Samples

We next examined SVs reported on chr1 of 5 individuals from the 1000 Genomes Project (11) to assess the sensitivity of VaPoR on real genomes (Table 1). We first observed that >95% of deletions and insertions could be successfully evaluated by VaPoR. For inversions, there were a limited number of events reported but at maximum only 1 event failed the VaPoR quality control per individual. A sensitivity of >90% was achieved for deletions (Figure 3a) and >80% for insertions (Figure 3b). To examine the false validation rate of VaPoR, we modified reported events on chr2 to appear at the same coordinates on chr1 and assessed them as though they were real events using the same sequence data set. VaPoR validated very few deletions or inversion and <10% of insertions.

**Figure 3.**
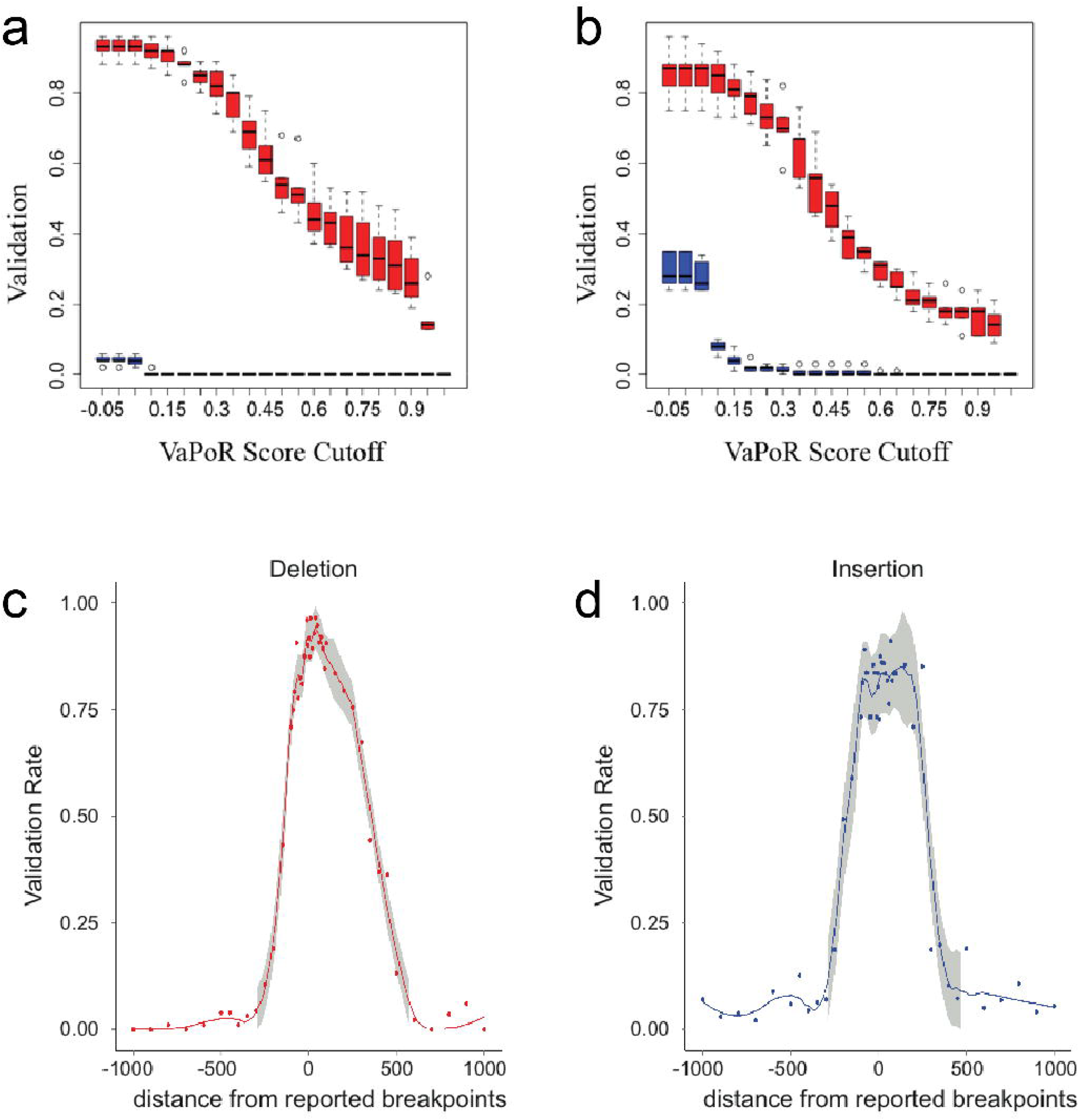
Validation rate and breakpoint accuracy of VaPoR on the 1000 Genomes Projects phase 3 calls. VaPoR was applied on 5 individuals with reported SVs as a truth set: HG00513, HG00731, HG00732, NA19238, NA19239. The validation rate of deletions (a) and insertions (b) are shown here across different cutoff scores for VaPoR. Robustness to breakpoint accuracy was assessed by deviating breakpoints from their actual positions across varying distances for deletions (c) and insertions (d).

We next assessed the performance of VaPoR to validate SVs with varying degrees of breakpoint accuracy. Real coordinates were artificially shifted each direction by −1000 to 1000 base pairs and reassessed with VaPoR for both simulated and real samples. In both cases, VaPoR exhibited a robust validation score up to approximately 200bp overall, with some slight differences observed between different SV types (Figure 3c,d).

### Discrimination of SV types and genotypes

We identified a small number of SVs in the high quality 1000 Genomes set that did not validate with VaPoR. Previous studies have shown that complex rearrangements are often misclassified as simple structural changes (3,12), and indeed upon manual inspection these appeared to consist of multiple connected rearrangements. For example, we observed a reported inversion in HG00513 and NA19239 on chromosome 1 (chr1:239952707-239953529) that was invalidated by VaPoR; an investigation into the long-reads aligned in the region showed the signature of an inverted duplication (Figure 4a) which, when incorporated into a modified reference that location, matched almost exactly with the read sequence (Figure 4b).

**Figure 4.**
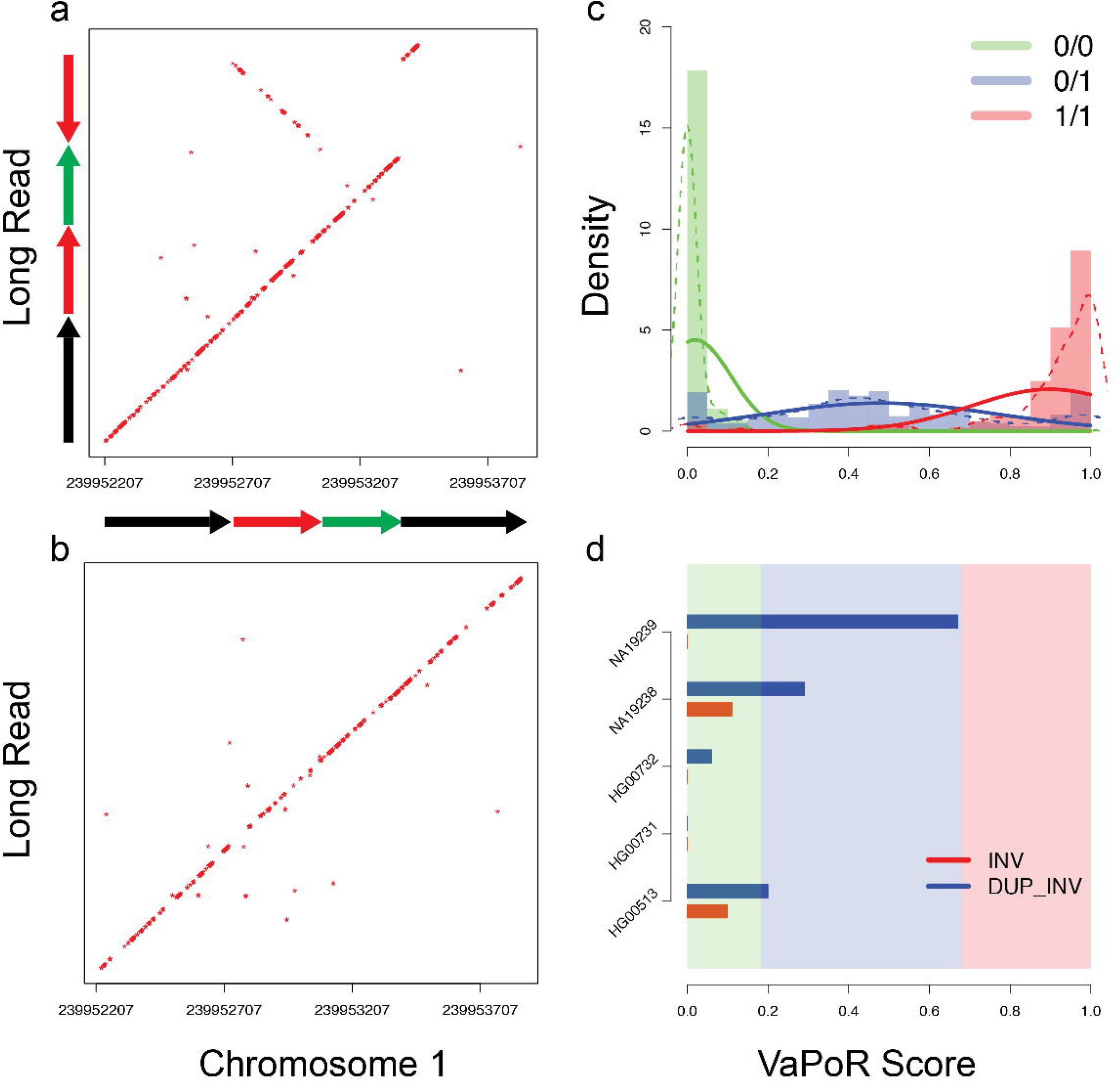
Validation and genotyping of assessed regions using VaPoR. (a) Recurrence plot of reference genome (GRCh38) to an aligned long read in NA19239 (m 150208_160301_42225_c100732022550000001823141405141504_s1_p0/3 831/0_12148) for a reported inversion at position chr1:239952707-239953529. The signature is consistent with an inverted duplication structure. (b) Recurrence plot of a different read (m150216_212941_42225_c100729442550000001823151505141565_s1_p0/106403/0_13205) against the same location, consistent with a non-variant (reference) structure. (c) Distribution of VaPoR scores on all reported SVs on chr1 in samples HG00513, HG00731, HG00732, NA19238, NA19239, stratified by color (solid) and modeled with a Guassian mixture model (dashed). (d) VaPoR scores of SV above now stratified by color as indicated in (c) for both reported inversion (red) and predicted inverted duplication (blue).

We further explored the distribution of VaPoR scores for this region and others across the sample set and observed clear delineations between allelic copy number when fit with a Gaussian mixture model allowing for the generation of genotype likelihoods for each site (Figure 4c). These tracked with our expected genotypes for the inverted duplication on chr1 across the 5 individuals queried while showing no support for the originally predicted inversion (Figure 4d). This shows that VaPoR is not only able to accurately genotype variants but can also distinguish between similar but distinct SV predictions in the same region.

### Runtime and efficiency

The computation runtime of VaPoR was assessed using 2 Intel Xeon Intel Xeon E7-4860 processors with 4GB RAM each on both simulated and real genomes. The runtime of simulated event was observed to increase linearly with read depth (Supplemental Figure 7). For events sequenced up to 20X, VaPoR takes ~3 seconds to assess a simple SV and ~5s for a complex event. The assessment of real samples sequenced at 20X required ~1.4 seconds to assess a simple deletion or insertion and ~6 seconds for an inversion (Supplemental Table 4).

## DISCUSSION

Here we present an automated assessment approach, named VaPoR, for exploring various features of predicted genomic structural variants using long read sequencing data. VaPoR directly compares the input reads with the reference sequences with relatively straightforward computational metrics, thus achieving high efficiency in both run time and computing cost. VaPoR exhibits high sensitivity and specificity in both simulated and real genomes, with the capability of discriminating partially resolved SVs either consisting of similar but incorrect SV types at the same location or correct SVs with offset breakpoints. Furthermore, we show that VaPoR performs well at low read depths (5-10X), thus providing the option of systematically assessing large-scale SVs with a lower sequencing cost.

## Availability and Requirements

Project name: VaPoR

Project home page: https://github.com/mills-lab/vapor

Operating systems: Linux, OS X

Programming languages: Python, R

Other requirements: Python v2.7.8+, rpy2, HTSeq, samtools v0.19+, pyfasta v0.5.2+, and pysam 0. 9.1.4+.

## Acknowledgements

We thank the Human Genome Structural Variation Consortium (HGSVC) for generating and providing the deep PacBio sequencing. We also thank Yuanfang Guan and Kerby Shedden for discussions over specific statistical considerations.

## Funding

This work was supported by the National Institutes of Health [R01HG007068]. AMW was supported by the Genome Science Training Program at the University of Michigan [T32HG000040]

## Authors’ contributions

XZ designed the algorithm, wrote the program, comparatively benchmarked the different algorithms, and wrote the manuscript. AMW generated simulated data, aided in assessment testing, and revised the manuscript. REM conceived the study, modified the algorithm, and revised the manuscript. All authors read and approved the final manuscript.

## Competing interests

None declared.

